# Direction of oscillatory networks in the human medial temporal lobe reverses across verbal working memory phases

**DOI:** 10.64898/2026.11.11.765952

**Authors:** Dongju Choi, Hio-Been Han

**Affiliations:** SeoulTech-KIRAMS Graduate School of Medical Sciences, Seoul National University of Science and Technology, Seoul 01811, Republic of Korea

## Abstract

Working memory (WM) requires neural circuits to change how they interact as information is encoded, maintained and used. Directional oscillatory interactions in the medial temporal lobe (MTL) contribute to memory processing, but how these interactions are spatially organized during verbal WM remains unclear. Here, we analyzed intracranial electroencephalogram and neuronal spiking recorded from the human MTL during a verbal Sternberg task. Theta- and gamma-band activities in MTL showed predominant directionality in the posterior-to-anterior axis during WM encoding, but this direction was reversed to anterior-to-posterior during retrieval. Theta activities reflecting the WM load were more prevalent in left than right MTL, and theta in left MTL coordinated spike timings more strongly. Moreover, interhemispheric interactions showed a predominant left-to-right influence, with posterior-to-anterior interactions enhanced during encoding and anterior-to-posterior interactions enhanced during retrieval. Together, these findings suggest that verbal WM engages a spatially organized MTL network in which the predominant direction of oscillatory interactions changes across task phases.

## Introduction

Working memory (WM) allows information to guide behavior after it is no longer present^1^. Doing so requires the same neural circuits to change how they interact as information is encoded, maintained and used. Recordings in the prefrontal cortex suggest that WM is carried by transient, oscillation-linked bursts of activity rather than by uniform persistent firing, and that the spatial organization of this activity contributes to WM control^2–4^. Theta rhythms are part of this picture: cortical theta increases throughout the Sternberg task^5^, scales with memory load^6^, and its phase in frontal cortex modulates WM readout^7^.

The direction of oscillatory interactions offers one way in which a network can change its mode of operation during memory processes. Theta and alpha oscillations travel across the human neocortex^8^, and during episodic memory their predominant direction is posterior-to-anterior at encoding and anterior-to-posterior at memory recall^9^, showing that the directional influence in this network changes upon task demands^10^. Hippocampal theta also propagates spatially in rodents and humans^11–14^, and the anterior and posterior hippocampus differ in connectivity and function^15,16^. Yet, whether interactions within the human medial temporal lobe (MTL) change direction in a comparable way across WM processes remains unclear.

The MTL contributes to WM. Hippocampal and other MTL neurons remain active during WM delays^17,18^, and hippocampal firing and hippocampal–cortical synchronization vary with verbal memory load^19^. Directional interactions involving the MTL have also been described in this verbal Sternberg paradigm. Granger causality between auditory cortex and hippocampus reverses between encoding and maintenance^20^. Within the MTL, posterior-to-anterior hippocampal theta/alpha influence accompanies correct maintenance^21^, information flows in both directions between amygdala and hippocampus during encoding and maintenance^22^, and hippocampal-to-entorhinal theta/alpha influence increases with load^23^. These studies focused on the maintenance period or compared it with encoding. How the predominant direction along the anterior–posterior axis evolves from encoding through maintenance to retrieval within a single trial during WM has not been described.

A second dimension of spatial organization of verbal WM is hemispheric. Verbal functions in the human brain preferentially engages left frontal and medial temporal regions^24–27^, and human hippocampal theta differs between hemispheres in its prevalence and frequency^28^ and in its relation to spatial memory^29^. In mice, hippocampus-dependent 5–7 Hz oscillations in the right cingulate–amygdala circuit are required for vicarious learning^30^, indicating that theta in mammalian limbic circuits can be recruited on one side according to the function involved. Whether verbal WM is accompanied by a leftward bias in human MTL, and whether interactions between the two MTLs are directional, has not been elucidated.

We addressed these questions using a human intracranial recording dataset of Boran et al.^19,31^, which contains intracranial electroencephalogram (iEEG) and single-neuron recordings from the hippocampus (Hipp), amygdala (Amg) and entorhinal cortex (EC) of nine patients performing a verbal Sternberg task. We characterized directional interactions between posterior hippocampus and more anterior MTL sites across the trial, theta and gamma bursts in each hemisphere, spike– field coherence, and interactions between the hemispheres. Theta- and gamma-band interactions were predominantly posterior-to-anterior during encoding and anterior-to-posterior during retrieval. Theta bursts were more frequent in the left MTL and reflected memory load there, left MTL neurons were more coherent with theta, and interhemispheric interactions ran predominantly from left to right, following the anterior–posterior direction of each task phase.

## Results

### Memory load slowed responses and reduced accuracy

Nine patients with drug-resistant epilepsy performed a modified Sternberg task (**Fig. 1a**)^19,31^. On each trial, a set of consonants (4, 6, or 8 letters) was shown for 2 s (encoding) and followed by a 3-s blank delay (maintenance). A probe letter then appeared, and the participant indicated whether it had been part of the set (retrieval). Reaction times (RT) increased and accuracy decreased with set size (**Fig. 1c**; reaction time, 1.23 ± 0.05, 1.46 ± 0.06 and 1.51 ± 0.07 s; accuracy, 98.8 ± 0.05, 91.2 ± 0.01 and 85.4 ± 0.02 % for set sizes 4, 6 and 8, respectively), as reported previously for this dataset^19^.

**Fig. 1.**
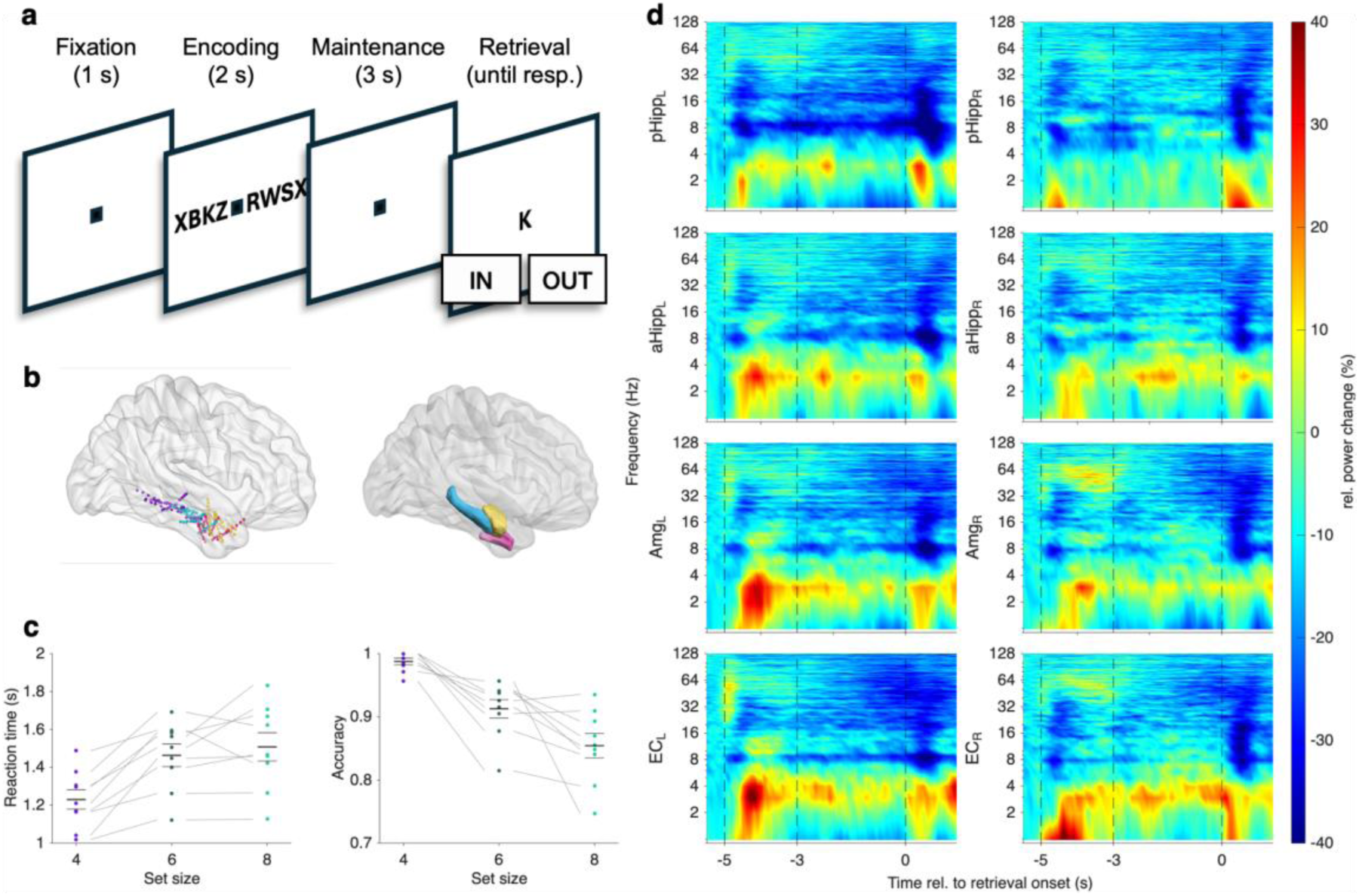
Task, recording sites, behavior and MTL power. **a**, Verbal Sternberg task. Fixation (1 s), encoding of four, six or eight letters (2 s), maintenance (3 s) and retrieval (probe; IN/OUT response). **b**, Contact locations in MNI space (sagittal, coronal and axial views): hippocampus (cyan), amygdala (yellow) and entorhinal cortex (magenta). **c**, Reaction time (left) and accuracy (right) as a function of set size. Dots, individual participants; black bars, mean ± s.e.m. **d**, Time– frequency power in pHipp, aHipp, Amg and EC in the left (left column) and right (right column) hemispheres, aligned to probe onset. Power is shown as the median percentage change across trials relative to a session-level fixation baseline, which is the median power across trials within each session, during the fixation period (n_sessions=37, n_trials=1541). Dashed lines mark encoding onset (−5 s), maintenance onset (−3 s) and probe onset (0 s).

We analysed iEEG from Hipp, Amg and EC contacts (**Fig. 1b**). Hipp contacts were labeled as posterior (pHipp) or anterior (aHipp) according to their position relative to the uncal apex (see **Methods**). In all regions, power below ~8 Hz increased after encoding onset and again after probe onset (**Fig. 1d**). Power between ~8 and 30 Hz decreased around retrieval, as alpha/beta power does in the prefrontal cortex when WM content is read out^32^.

### Oscillatory interactions reversed direction between encoding and retrieval

To characterize the direction of interactions along the anterior–posterior axis, we computed spectrally and temporally resolved Granger causality (GC) for each ipsilateral pair linking a pHipp to a more anterior contacts (i.e., aHipp, amygdala or EC). Total 42 pairs from 8 patients. One patient who didn’t have any ipsilateral pairs is excluded. Net GC (i.e., *bias-to-anterior*) was defined as GC from the posterior to the anterior site minus GC in the opposite direction, so that positive values indicate a predominantly from-posterior-to-anterior interaction (see **Methods**).

The predominant direction depended on the task phase (**Fig. 2**). In the example pair in **Fig. 2c**, GC from pHipp to EC was elevated during encoding, whereas GC from EC to pHipp was elevated during retrieval. Across pairs, net GC in theta-band (4–10 Hz) became positive shortly after encoding onset and remained positive into the early delay (**Fig. 2d**). It became negative in the second half of the delay, about 2 s before probe onset, and remained negative throughout the retrieval period.

**Fig. 2.**
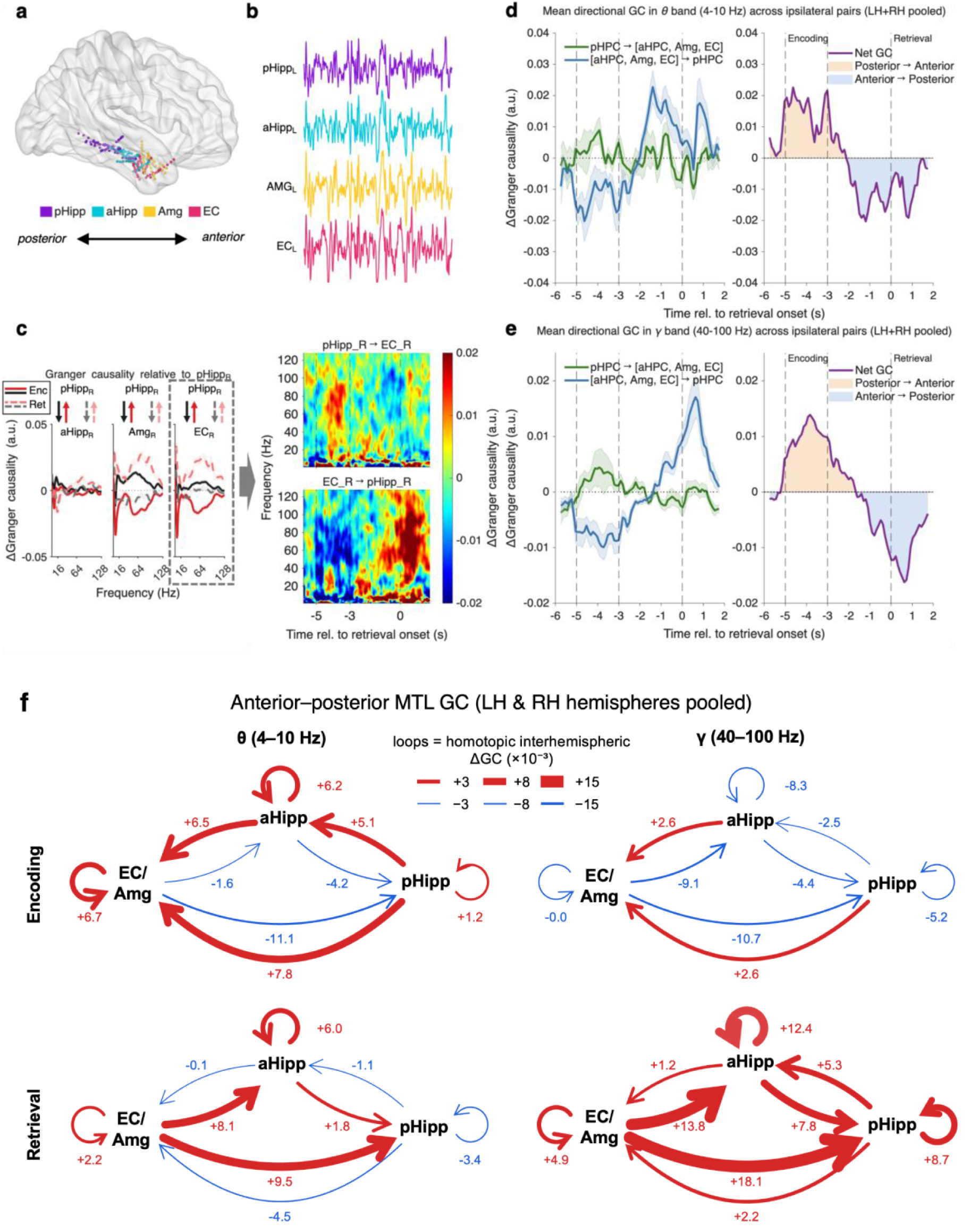
Theta- and gamma-band interactions reversed direction between encoding and retrieval. **a**, Schematic of the long axis, with recording sites from posterior hippocampus to anterior MTL. **b**, Example raw signals from right pHipp, aHipp, Amg and EC. **c**, Left, spectral net GC relative to right pHipp for aHipp, Amg and EC, during encoding (solid) and retrieval (dashed). Right, time–frequency GC for pHipp→EC (top) and EC→pHipp (bottom) for an example right-hemisphere pair. **d**, Left, mean directional GC measure in the theta band (4–10 Hz) for posterior-to-anterior (green) and anterior-to-posterior (blue) directions across ipsilateral pairs (pair n = 42, trial n = 1541). Right, net GC; orange shading, posterior-to-anterior dominance; blue shading, anterior-to-posterior dominance. **e**, As in d, for the gamma band (40–100 Hz). Shading, ± s.e.m. across pairs. **f**, Directional interactions along the anterior–posterior axis of the MTL. Nodes represent EC/Amg (EC and Amg merged), aHipp and pHipp, with both hemispheres and both ipsilateral and contralateral pairs pooled. Each arrow shows the mean ΔGC (×10^−3^) from the source region to the target region during encoding (−5 to −3 s; top) and retrieval (0 to 1 s; bottom) relative to retrieval onset. Red, thick arrows denote increases and blue, thin arrows decreases. Self-loops denote homotopic interhemispheric interactions (for example, aHipp_L ↔ aHipp_R).

Gamma-band (40–100 Hz) interactions followed the same pattern (**Fig. 2e**). Net GC was positive during encoding (i.e., predominantly posterior-to-anterior), peaking about 1 s after encoding onset, and negative from the late delay onward, with the largest anterior-to-posterior bias about 1 s after probe onset.

### Theta bursts were more frequent in the left MTL and reflected memory load

Oscillatory activity during WM tends to occur in brief bursts^2^. We identified theta and gamma bursts at each brain region as periods in which the band-limited amplitude exceeded the 80th percentile for at least two cycles (**Fig. 3a**; Methods).

**Fig. 3.**
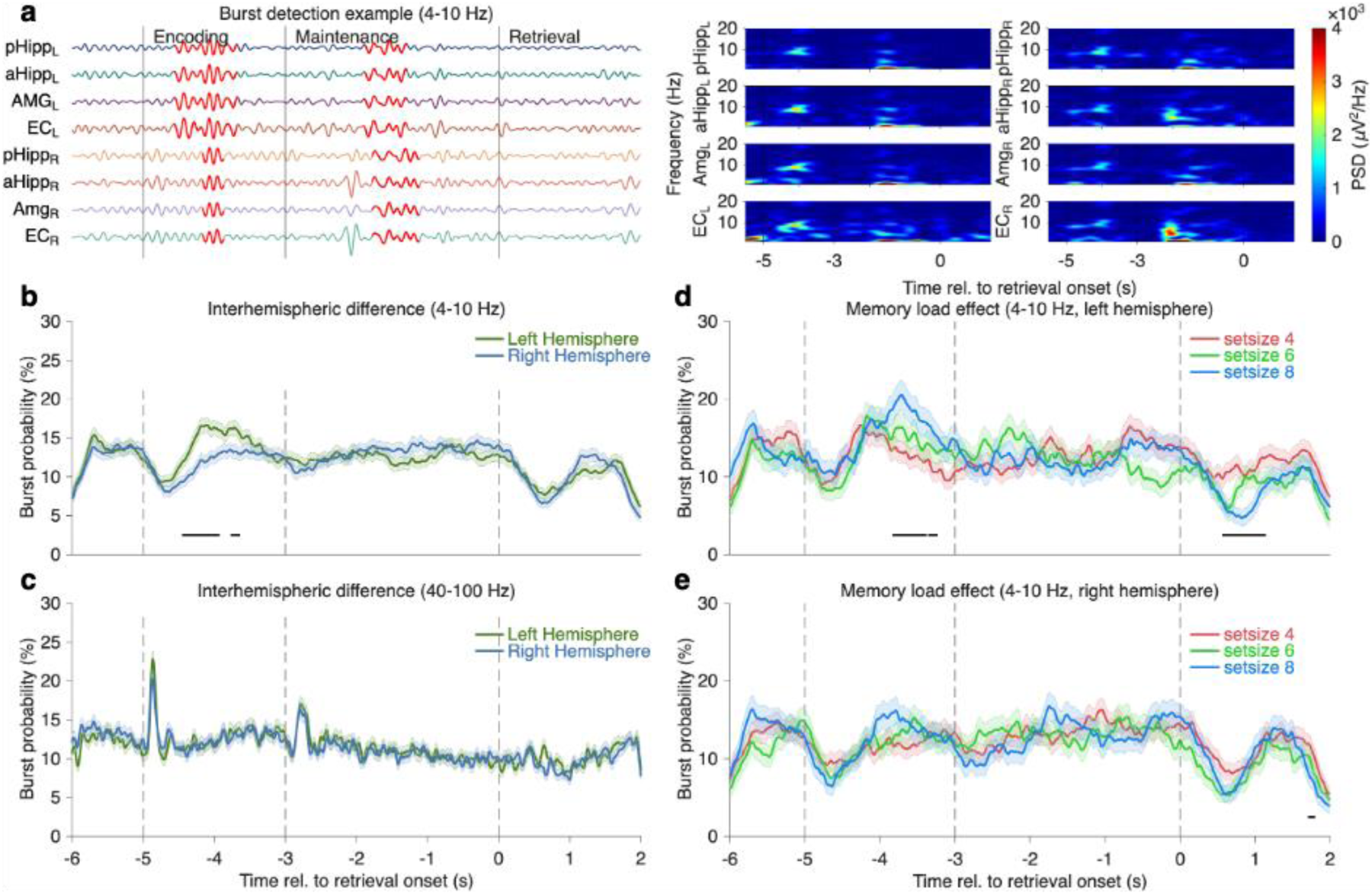
Theta bursts were more frequent in the left MTL and reflected memory load. **a**, Example of burst detection (4–10 Hz) across left and right MTL contacts during encoding, maintenance and retrieval. A burst was defined as an envelope above the 80th percentile lasting more than two cycles. **b**, Theta burst probability in the left (green) and right (blue) hemispheres. **c**, As in **b**, for gamma (40–100 Hz) bursts; no hemispheric difference. d, Left-hemisphere theta burst probability by set size (red, 4; green, 6; blue, 8). **e**, As in **d**, for the right hemisphere. Left panels, aligned to encoding; right panels, aligned to probe onset. Black bars, significant clusters (p < 0.05). Shading, ± s.e.m.

During encoding, theta bursts were more frequent in the left than in the right MTL (**Fig. 3b**). Unlike theta, bursts in the gamma band did not differ between hemispheres (**Fig. 3c**). In the left hemisphere, theta burst during encoding increased with set size and was highest for eight-letter sets (**Fig. 3d**). This load effect was not observed in the right hemisphere (**Fig. 3e**).

### Left MTL neurons fired in closer coordination with theta

We next asked whether local spike timing was coordinated with theta differently in the two hemispheres. For each unit, we computed spike–field coherence with the iEEG recorded in the same region (**Methods**). In pHipp, coherence at ~5–6 Hz was higher for left-hemisphere units (n = 66) than for right-hemisphere units (n = 43) during both encoding and retrieval (**Fig. 4a-b**; *p*s < .05). Likewise, left-hemisphere EC units (n = 111) showed a pronounced coherence peak in the theta range that was absent in right-hemisphere EC units (n = 201), again in both periods (**Fig. 4c-d**; *p*s < .05).

**Fig. 4.**
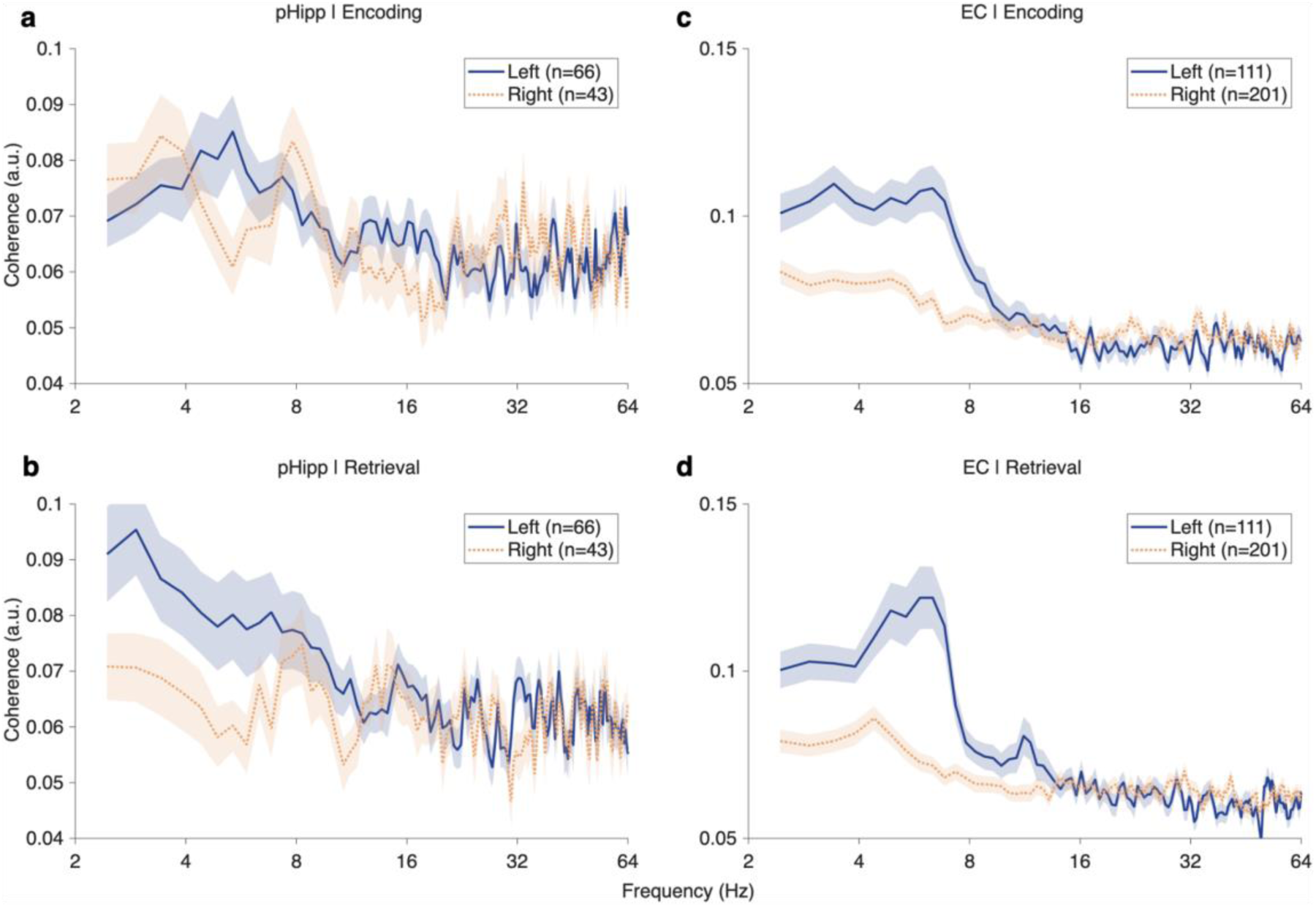
Left-hemisphere MTL neurons fired in closer coordination with theta. Spike–field coherence for units in pHipp (**a,b**; left, n = 66; right, n = 43) and EC (**c,d**; left, n = 111; right, n = 201) during encoding (**a,c**) and retrieval (**b,d**). Solid blue, left hemisphere; dotted orange, right hemisphere. Shading, ± s.e.m..

### Interhemispheric interactions were predominantly left-to-right

In patients with electrodes in both hemispheres (total 84 pairs – 42 ipsilateral pairs, 42 bilateral pairs – from 9 patients), we computed GC between contralateral MTL contacts. Interhemispheric interactions also followed the anterior–posterior direction of each task phase (**Fig. 5a**). From left posterior to right anterior sites, GC increased between ~20 and 60 Hz during encoding. From left anterior to right posterior sites, the largest increase occurred after probe onset, between ~30 and 80 Hz.

**Fig. 5.**
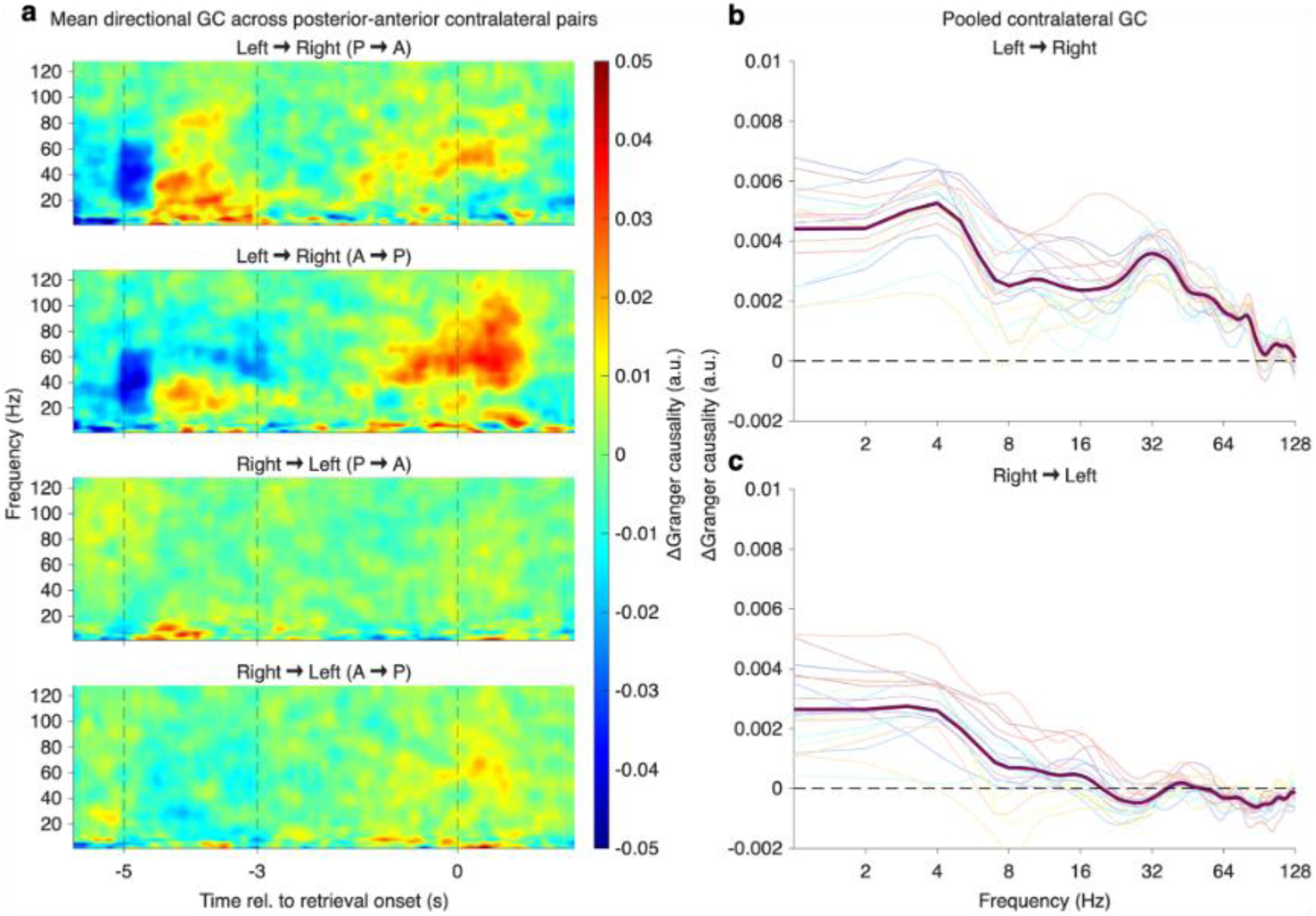
Interhemispheric interactions were predominantly left-to-right. **a**, Time–frequency net GC for contralateral pairs: left→right posterior-to-anterior, left→right anterior-to-posterior, right→left posterior-to-anterior and right→left anterior-to-posterior (top to bottom), aligned to probe onset. **b,c**, Spectral GC for left→right (**b**) and right→left (**c**) influence, averaged across the trial.

**Fig. 6.**
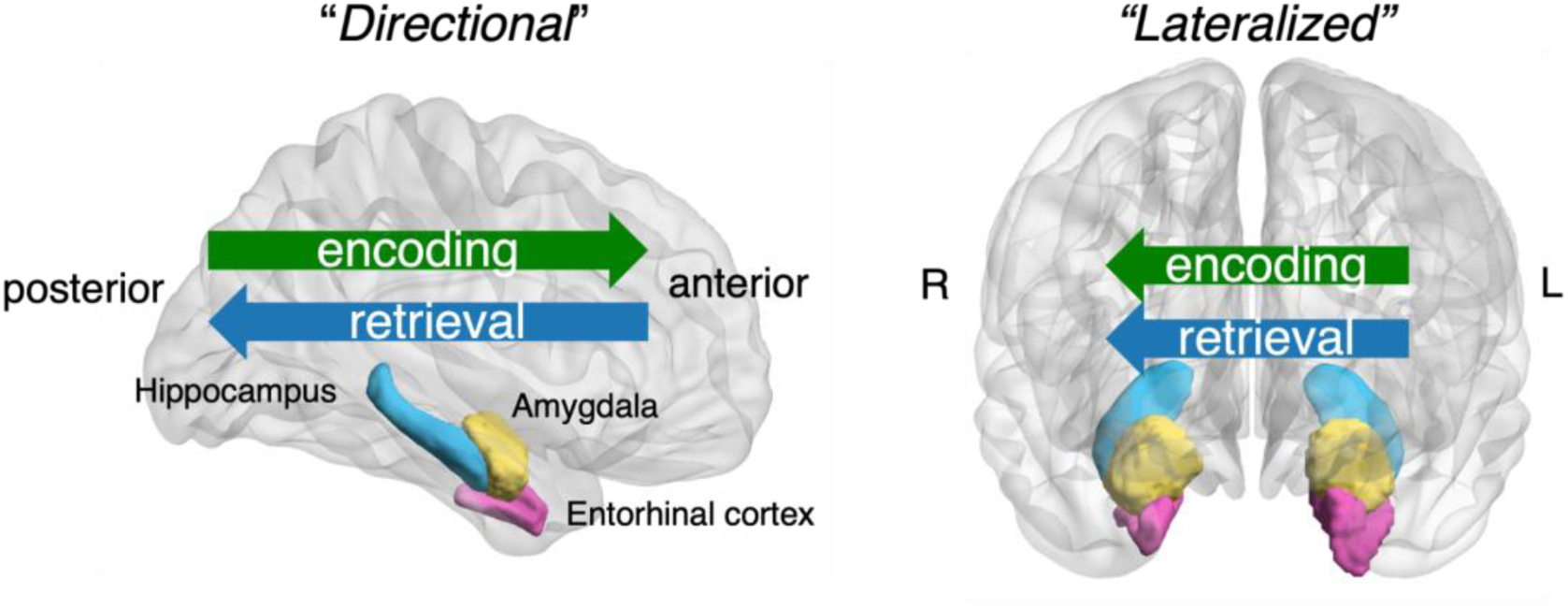
Summary. Verbal WM engages a spatially organized MTL network. Along the anterior–posterior axis, the predominant direction of oscillatory interactions is posterior-to-anterior during encoding and anterior-to-posterior during retrieval. Between hemispheres, theta bursts and spike–field coherence are stronger on the left, and interactions run predominantly from left to right. Arrows denote directed statistical interactions, not demonstrated wave propagation or anatomical transmission.

Averaged over the trial, net GC from left to right was above zero from 2 Hz to ~64 Hz and was largest near 4 Hz (**Fig. 5b**). GC from right to left was smaller throughout almost every other frequency band (**Fig. 5c**).

## Discussion

Verbal WM was accompanied by a spatially organized pattern of MTL interactions whose predominant direction changed across task phases. Theta- and gamma-band interactions between posterior hippocampus and anterior MTL sites were predominantly posterior-to-anterior during encoding and anterior-to-posterior during retrieval, with the change beginning in the late delay. Theta was biased toward the left hemisphere, where bursts were more frequent and varies with memory load, and where spikings were more coherent with theta. Interactions between the hemispheres ran predominantly from left to right as long as followed the anterior–posterior direction of each task phase. How organization along these two spatial dimensions is linked mechanistically remains to be tested.

Earlier studies in the same paradigm provide the closest comparison. Li et al. found that correct maintenance was associated with predominantly posterior-to-anterior theta/alpha influence within the hippocampus^21^, that hippocampal influence on the EC increased with load^23^, and that information flowed in both directions between amygdala and hippocampus^22^. Our early-delay results agree with the posterior-to-anterior direction, but the late delay showed the opposite direction, which appears to differ from their maintenance-averaged result. The analyses differ in temporal resolution, frequency range, trial selection and included region pairs: our pooled contrast also contains amygdala and EC sites. Reproducing the earlier analysis in pHipp–aHipp pairs before separating early and late delay would show whether the two sets of results reflect different stages of the same within-trial time course.

The reversal resembles the change in direction of cortical theta and alpha traveling waves between episodic encoding and recall^9^. Here a comparable change occurred within the MTL and within a single trial lasting about 6 s. GC measures directed predictability between sampled sites, whereas traveling-wave analyses characterize spatial phase progression, and amygdala and EC contacts lie outside the hippocampal long axis. The present data therefore do not establish a continuous wave, its speed or its geometry. They do indicate that the predominant direction of interaction is a task-dependent property shared by the MTL and the neocortex. Hippocampal– scalp theta–alpha coupling has been reported to be increase with memory load in this paradigm^19^, and hippocampal–auditory cortex interactions reverse between encoding and maintenance^20^; simultaneous MTL and cortical recordings could show whether the two change direction together.

The posterior–anterior axis of the human hippocampus corresponds to the septotemporal (dorsoventral) axis in rodents^33^. Gene expression, connectivity and function change gradually along this axis: the septal pole carries fine-grained spatial representations, whereas the temporal pole represents space at coarser scales and is preferentially connected with the amygdala, hypothalamus and medial prefrontal cortex^34–37^. Rodent theta travels from the septal to the temporal pole (i.e., septic-temporal axis)^38,39^, which corresponds to the posterior-to-anterior direction that predominated here during encoding, and theta at the temporal pole is weaker and less coherent with septal theta^39,40^. Such regional dissociation of theta is not unique to the hippocampus: in the mouse neocortex, frontal and visual theta are modulated differently during sustained attention^41^. In mice, direct ventral hippocampal input to the medial prefrontal cortex is required for encoding, but not for maintenance or retrieval, of spatial WM and is accompanied by gamma-band synchrony between the two structures^42^, and prefrontal theta synchronizes with the hippocampus through its ventral part^43^. Oscillatory interactions between the prefrontal cortex and the amygdala also switch within seconds according to behavioral demands in mice^44^. One possibility is therefore that posterior-to-anterior interaction during encoding conveys item information toward the anterior output stage of the hippocampus, whereas the reversal toward retrieval reflects the influence of anterior MTL sites (e.g., EC), which are positioned to relay prefrontal and amygdalar signals, on the posterior hippocampus. In humans, frontal theta directs MTL activity during WM processing^45^. This account remains to be tested: the rodent evidence comes from spatial tasks, and the present recordings did not include the prefrontal recordings.

The change in direction began while the screen was blank, so it cannot be solely attributed to the probe. It may reflect internal rehearsal, which participants report in this task^19^ and which is accompanied by hippocampal-to-cortical influence^20^, preparation to compare the probe with the remembered letters, or anticipation of a predictable event. Because the delay had a fixed duration, these possibilities cannot be separated here. Tasks with variable delays, and analyses relating the timing of the change to behavior, would distinguish them. Models in which encoding and retrieval occupy different phases of the theta cycle^46^ suggest a further test: whether the two directional states are also associated with different theta phases of local spiking.

The findings are consistent with accounts in which WM depends on transient, oscillation-linked activity whose spatial organization supports control^2–4^. MTL theta occurred in bursts, and in the left MTL their probability reflected memory load, as gamma burst rates do in prefrontal cortex^2^. The decrease in 8–30 Hz power around retrieval parallels the drop in prefrontal beta bursts when WM content is read out^32^. The present data do not test item-specific gamma coding, the movement of item representations or the absence of persistent firing. Relating the directional state to burst timing, spike coupling or behavior on single trials would connect the findings more directly to these accounts.

The leftward bias in theta bursts and spike–field coherence is consistent with the involvement of the left MTL in verbal memory^24–27^ and with the higher prevalence of left hippocampal theta bouts reported during recognition memory^28^. Lateralization of limbic theta is not fixed to the left: in rodents, right-hemisphere cingulate–amygdala theta supports observational fear^30^. Which hemisphere is engaged may therefore depend on the function. Testing this in humans would require a matched nonverbal or affective condition, which the present task lacks. The coherence difference also needs to be assessed with spike counts, trial counts and patient contributions controlled. Direct commissural connections between the human hippocampi are limited^47^, so the left-to-right interactions are probably indirect.

Our study has some limitations. Most of all, GC is sensitive to signal quality, reference choice, common inputs and nonstationarity^48^. Showing that the directional state predicts behavior would strengthen its functional interpretation. A causal test could manipulate the relative timing of posterior and anterior hippocampal activity; temporal interference stimulation offers a possible route to the human hippocampus^49,50^. Such a test would require independent validation of the induced spatial phase pattern and would not follow from GC direction alone.

## Methods

### Dataset and participants

We analysed the publicly available dataset of Boran et al^19,31^. Nine patients with drug-resistant focal epilepsy (5 female; age range 18–56 years; all right-handed) were implanted with depth electrodes for seizure localization^19,31^. Electrode placement was determined solely by clinical criteria. The original study was approved by Kantonale Ethikkommission Zürich, PB-2016-02055^19^.

### Task

Participants performed a modified Sternberg task (**Fig. 1a**)^19^. Each trial began with a 1-s fixation period. A string of eight letters was then presented for 2 s (encoding); the central four, six or eight letters were consonants to be memorized (set size) and the remaining outer positions were filled with “X”, which was never a memory item. A probe letter then appeared, and participants pressed one of two buttons to indicate whether it had been part of the set (IN/OUT). Set sizes were presented in random order, except that a trial with an incorrect response was always followed by a set-size-4 trial. Each session comprised 50 trials, and patients completed two to seven sessions (37 sessions in total).

### Recordings and electrode localization

The iEEG data were recorded from depth electrodes (1.3 mm diameter, eight contacts, 5 mm spacing; AD-Tech), and unit activity from nine microelectrodes protruding ~4 mm from each electrode tip, against a common intracranial reference at 4 kHz and 32 kHz, respectively (ATLAS, Neuralynx)^31^. The iEEG provided with the dataset was resampled to 2 kHz. Contacts were assigned to the hippocampus, Amg or EC using the MNI coordinates provided with the dataset30. `Hippocampal contacts anterior to the uncal apex (MNI y = −21 mm) were labelled aHipp and those posterior to it pHipp`^16^. In total, 271 contacts on 34 electrodes in the left MTL and 215 contacts on 27 electrodes in the right MTL were used.

### Preprocessing

Signals from the same brain area label were averaged to produce region-representative iEEG signals (e.g., pHipp, aHipp, Amg, and EC in each hemisphere, total 8 region-representative iEEG signals). Bad contacts were identified separately for each session using two criteria computed over all trials. First, contacts whose signal standard deviation deviated by more than 1.69 s.d. from the mean across all contacts were excluded. Second, among the remaining contacts of each electrode, those whose signal correlated weakly (Pearson r < 0.5) with the median signal of that electrode were excluded. The remaining contacts were averaged to produce one region-representative iEEG signal per region and hemisphere (pHipp, aHipp, Amg and EC; eight signals in total). Trials marked as containing artefacts in the dataset were excluded. `Data were aligned to probe onset (−6 to +2 s), with fixation from −6 to −5 s, encoding from −5 to −3 s and maintenance from −3 to 0 s.`

### Spectral power

Time–frequency power from 1 to 128 Hz was estimated with fast Fourier transform. Band-limited power change was expressed as median percentage change across trials from session-level baseline, which is the median of trial-wise power during the fixation period.

### Granger causality

Directed interactions were assessed with time- and frequency-resolved GC^51,52^ computed in sliding windows (0.5 s length, 200 ms step, model order is set to 100 ms). For each long-axis pair, the net directional contrast (i.e., *bias-to-anterior*) was defined as follow; ΔGC(f, t) = GC posterior→anterior(f, t) − GC anterior→posterior(f, t).

Long-axis pairs linked pHipp to aHipp, amygdala or EC in the same hemisphere. For interhemispheric analyses, net GC was defined as left-to-right minus right-to-left GC, separately for contralateral posterior-to-anterior and anterior-to-posterior pairs. Band-limited time courses were obtained by averaging over 4–10 Hz (theta) and 40–100 Hz (gamma). The theta band was chosen to include both the slower and faster theta rhythms described in the human hippocampus^53,54^. A comparable time-resolved GC approach has been used to track bidirectional frontoparietal interactions during WM^55^.

### Burst detection

Signals were band-pass filtered in the theta (4–10 Hz) and gamma (40–100 Hz) bands, and the amplitude envelope was obtained with the Hilbert transform. A burst was defined as an epoch in which the envelope exceeded the 80th percentile of that contact’s amplitude distribution, computed over the whole session for each trial, for at least two cycles of the band’s centre frequency^56^. Burst probability was the proportion of trials with a burst at each time point, averaged across contacts within each hemisphere.

### Spike–field coherence

Spikes were detected and sorted as described previously^19,31^. For each unit, coherence was computed between its spike train and the field potential of a macro-contact in the same region and hemisphere. Coherence was computed separately for the encoding window and the retrieval window from 2 to 64 Hz using Chronux toolbox^57^.

### Statistics

Behavioral measures, spectral power, GC and spike–field coherence were compared with non-parametric versions of t-tests (signed-rank or rank-sum) tests. For burst probability, a bootstrapping method was used. Burst probability was compared between hemispheres and between set sizes using cluster-based permutation tests with generalized linear mixed-effects models (GLMEs) with 1,000 permutations^58^.

## Code availability

Analysis code (MATLAB) will be made available upon request to the corresponding author.

## Acknowledgements

We thank Johannes Sarnthein and colleagues for making their dataset openly available. This work was supported by a research program funded by Seoul National University of Science and Technology (SeoulTech).

## Author contributions

D.C. and H.-B.H. conceived the study. D.C. analysed the data. H.-B.H. supervised the project. D.C. and H.-B.H. wrote the manuscript.

